# Cell overgrowth during G1 arrest triggers an osmotic stress response and chronic p38 activation to promote cell cycle exit

**DOI:** 10.1101/2022.09.08.506843

**Authors:** Lisa Crozier, Reece Foy, Rozita Adib, Mihaly Badonyi, Ananya Kar, Jordan A. Holt, Rona Wilson, Clement Regnault, Phil Whitfield, Joseph A. Marsh, Adrian Saurin, Alexis R. Barr, Tony Ly

## Abstract

Cell size and the cell cycle are intrinsically coupled and abnormal increases in cell size are associated with senescence. The mechanism by which overgrowth primes cells to exit the cell cycle remains unclear. We investigate this using CDK4/6 inhibitors that arrest cell cycle progression in G0/G1 and are used to treat ER+/HER2-metastatic breast cancer. We demonstrate that long-term CDK4/6 inhibition promotes cellular overgrowth during the G0/G1 arrest, causing widespread proteome remodeling and p38-p53-p21-dependent cell cycle exit. Cell cycle exit is triggered by two waves of p21 induction. First, overgrowth during a G0/G1 arrest induces an osmotic stress response, producing the first wave of p21 induction. Second, when CDK4/6 inhibitors are removed, a fraction of cells escape G0/G1 arrest and enter S-phase where overgrowth-driven replication stress results in a second wave of p21 induction that causes cell cycle withdrawal from G2, or the subsequent G1. This could explain why cellular hypertrophy is associated with senescence and why CDK4/6 inhibitors have long-lasting anti-proliferative effects in patients.

## INTRODUCTION

The success of CDK4/6 inhibitors (CDK4/6i) in treating estrogen-receptor positive (ER+), human epidermal receptor growth factor 2 negative (HER2-) metastatic breast cancer (Cristofanilli et al., 2016; Dickler et al., 2017; Im et al., 2019) has reinvigorated interest in targeting the pathways regulating cell cycle entry for cancer treatment (Álvarez-Fernández and Malumbres, 2020; Fassl et al., 2022; Goel et al., 2022; Matthews et al., 2022). CDK4/6 forms active kinase complexes with D-type cyclins that phosphorylate the retinoblastoma protein (Rb) (Matsushime et al., 1992) to promote G0-G1 progression and S-phase entry (Sherr, 1993). Inhibition of CDK4/6 results in hypophosphorylated Rb, which suppresses transcription of cell cycle genes. Results from the clinic, animal models, and *in vitro* suggest that the mechanisms of action of CDK4/6i are complex (Knudsen and Witkiewicz, 2017). For example, while the high frequency of resistance-driving genetic lesions in RB1 was expected, the mechanisms of how other mutations drive CDK4/6i resistance cannot be so easily explained (Gong et al., 2017; Wander et al., 2020). CDK4/6i has unanticipated effects on the immune system (Deng et al., 2018; Goel et al., 2017; Heckler et al., 2021; Lelliott et al., 2021), and tumour cell clearance in animal models of Her2+ breast cancer is dependent on T-cell lymphocyte function (Goel et al., 2017). Thus, major gaps exist in our current understanding of the molecular mechanisms of CDK4/6i-mediated inhibition of tumour cell proliferation and tumour cell clearance.

Previous work has shown that prolonged treatment with CDK4/6i can promote senescence across a range of cell types (Barr and McClelland, 2022; Crozier et al., 2022; Lengefeld et al., 2022; Neurohr et al., 2019; Wagner and Gil, 2020; Wang et al., 2022). The source of the stress that leads to cell cycle exit remains unclear, although recently this has been linked to replication stress (Crozier et al., 2022). CDK4/6i treated cells arrest in G0/G1 but continue to accumulate biomass and increase in cell volume (Lengefeld et al., 2022; Neurohr et al., 2019; Tan et al., 2021). Continued cell growth contributes to the onset of permanent cell cycle arrest and senescent phenotypes (Korotchkina et al., 2010), however, the mechanisms linking excessive cellular growth to permanent cell cycle arrest are poorly characterized. A recent study proposed a model whereby excessive growth produces a diluted cytoplasm that limits cellular functions (Neurohr et al., 2019). Downstream, permanent cell cycle arrest after prolonged CDK4/6i treatment has been demonstrated to be p53-dependent (Crozier et al., 2022; Manohar et al., 2022; Wang et al., 2022), but the molecular pathways that link excessive cell growth to p53 activation are not understood.

Cell cycle exit is correlated with the length of time cells spend in CDK4/6i (Crozier et al., 2022; Trotter and Hagan, 2022). Here, and in the accompanying paper (Foy et al., 2022), we demonstrate that permanent cell cycle arrest is caused by excessive cell growth during G1 arrest and not treatment duration *per se*. We show that prolonged CDK4/6i treatment promotes cell overgrowth, a cell state characterized by abnormally large size, cell stress, p53 activation and increased levels of the CDK inhibitor, p21. This causes a fraction of cells to exit from the cell cycle in G0/G1. The remaining cells enter S-phase when CDK4/6i is removed, and experience overgrowth-dependent replication stress, which promotes a second wave of p21 expression after S-phase completion. This second p21 wave elicits cell cycle withdrawal from G2, or from the subsequent G1. Our work establishes how cellular overgrowth can be sensed by cells to promote cell cycle exit.

## RESULTS

### Long-term CDK4/6 inhibitor treatment promotes cellular overgrowth that drives permanent cell cycle arrest

Treatment of hTERT-RPE1 (RPE1) cells with the CDK4/6 inhibitor, palbociclib (CDK4/6i), causes cells to arrest in G0/G1 (Trotter and Hagan, 2022). Prolonged treatment of RPE1 cells with CDK4/6i for up to seven days was accompanied by an increase in cell size that correlated with the length of CDK4/6i treatment (**Figures 1A, B; Supplementary Figure 1**). This increase in cell size was accompanied by increases in the cellular protein and RNA content (**Figures 1C, D**). It has been previously reported that the ability of cells to re-enter S-phase after CDK4/6i washout starts to decline after two days of CDK4/6i treatment (Crozier et al., 2022; Trotter and Hagan, 2022). We observed that this ability to re-enter S-phase after CDK4/6i treatment negatively correlates with treatment duration and cell size (**Figure 1E**).

**Figure 1.**
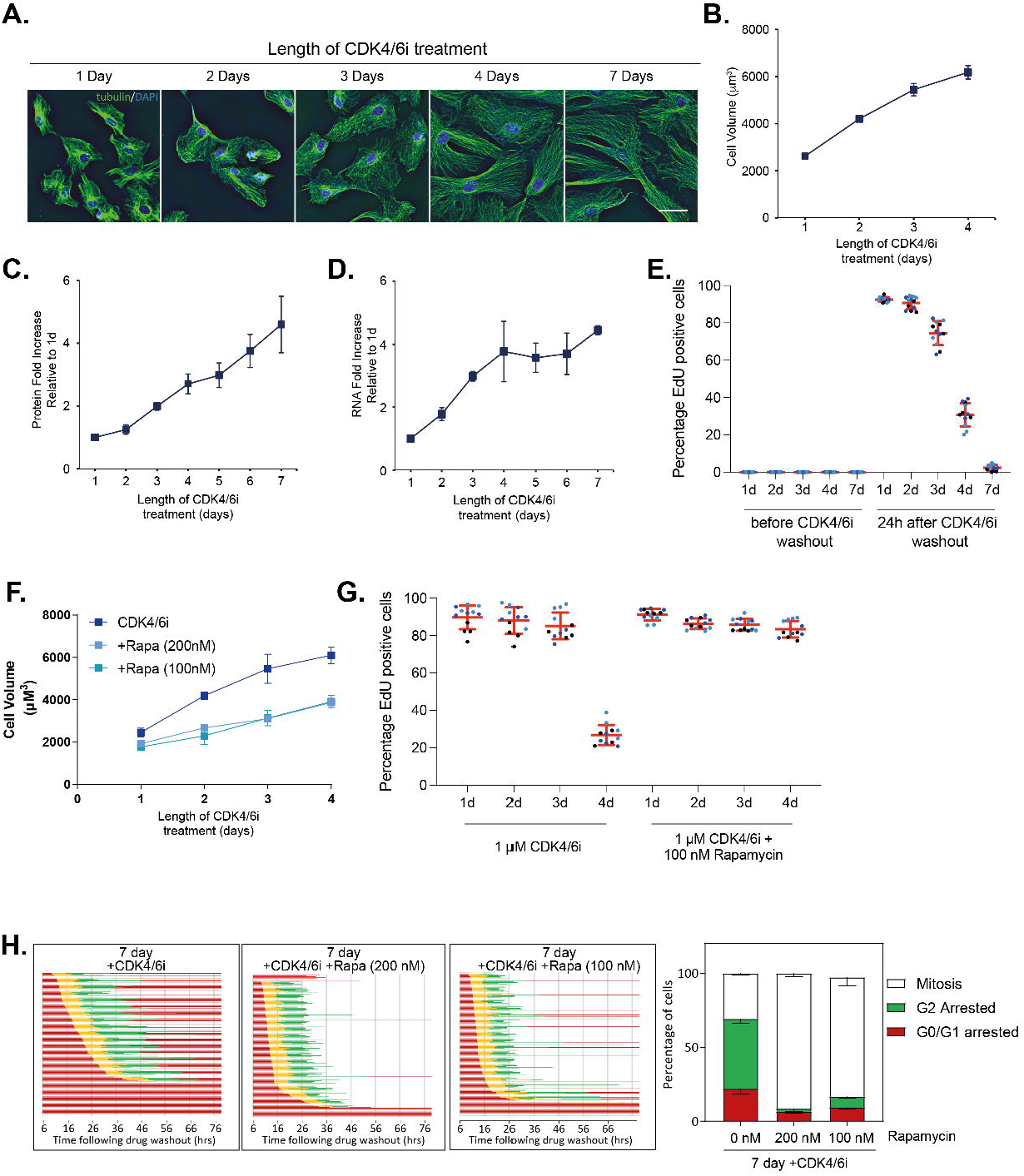
Long-term CDK4/6 inhibitor treatment promotes cellular overgrowth that drives permanent cell cycle arrest. **A**. Images of RPE1 cells incubated in CDK4/6i for increasing time. In green is alpha-tubulin to label the cytoplasm, blue is DAPI to label nuclei. Scale bar is 50 um. **B-D**. Cells continue to grow during a prolonged CDK4/6i arrest shown by increases in cell volume (**B**), protein content (**C**) and RNA content (**D**). Mean of n=2 +/- s.d. are shown. Note, in (**B**) cell volume is only measured up to 4d of CDK4/6i treatment because CDK4/6i-treated RPE1 cells do not round up properly after trypsinisation beyond 4d making accurate cell volume measurements challenging. **E**. Graph showing percentage of cells re-entering S-phase (EdU positive) after increasing days of CDK4/6i treatment followed by washout. RPE1 mRuby-PCNA cells were arrested in 1 μM CDK4/6i for the indicated times, inhibitor was then washed out and fresh media containing EdU was added. Cells were fixed after 24h to quantify cells that had re-entered S-phase. Data are plotted as superplots where each technical repeat is a single dot and biological repeats are in different colours. Mean +/- s.d. of n=3 are shown in red. **F**. Co-treatment of cells with CDK4/6i and rapamycin inhibits cell growth during the prolonged arrest. Mean of n=3 +/- s.d. are shown. **G**. Co-treatment of cells with CDK4/6i and rapamycin allows cells to maintain proliferative potential during the prolonged arrest. Mean +/- s.d. of n=3 are shown in red. **H**. Graphs show FUCCI RPE1 cells released from 7 d arrest with indicated inhibitors into the Eg5 inhibitor STLC to arrest cells in mitosis. Live cell imaging was started 6h after drug washout. Red is G1, yellow is S-phase, green is G2. Cells that withdraw from the cell cycle in G2 return to red colour as APC/C^Cdh1^ becomes active and Geminin is degraded. Right-hand graph shows mean of n=3 +/- s.d.

To investigate if the increase in cell size causes this reduced ability to re-enter S-phase after CDK4/6i washout, we treated cells with CDK4/6i under conditions that restricted cell growth. mTOR is a master regulator of cell growth and co-treatment with rapamycin, an inhibitor of mTORC1 activity, reduced cell growth during CDK4/6i treatment (**Figure 1F**). Indeed, co-treatment of cells with CDK4/6i and rapamycin restored the ability of cells to re-enter S-phase after inhibitor washout, suggesting that cell overgrowth, and not CDK4/6i treatment duration *per se*, is the main driver of long-term G0/G1 arrest after CDK4/6i washout (**Figure 1G**). To better understand cell cycle behaviour after CDK4/6i washout, we used live cell imaging of RPE1 FUCCI cells (Krenning et al., 2014), treated with either CDK4/6i alone, or combined CDK4/6i and rapamycin (**Figure 1H**). 22% of CDK4/6i-treated cells remain in a G0/G1 arrest after CDK4/6i washout, and of those cells that did re-enter S-phase, a large fraction withdrew from the cell cycle with 4N DNA content, as previously observed (Crozier et al., 2022). Preventing cell growth with rapamycin co-treatment allowed more cells to re-enter S-phase, with a shorter G1 length and a higher fraction of cells completing G2 and entering mitosis (**Figure 1H**).

Based on the results here and the accompanying paper by (Foy et al., 2022), we conclude that continued cellular growth induced by a prolonged G0/G1 arrest drives cell cycle exit following CDK4/6i treatment. To distinguish this from physiological growth during an unperturbed G1, we suggest that prolonged G1 arrest promotes *overgrowth*. We next investigated the molecular mechanisms driving the pathologies observed with cell overgrowth.

### Long-term cell cycle arrest induced by cellular overgrowth is p21-dependent

We previously showed that p21 protein starts to increase after two days CDK4/6i treatment (Pennycook and Barr, 2022) and p53, a key transcription factor for p21, was shown to be required for long-term cell cycle arrest after washout of prolonged CDK4/6i treatment (Crozier et al., 2022; Wang et al., 2022). The increase in p21 protein correlates with the loss of the ability of cells to re-enter S-phase after CDK4/6i washout between two and three days of CDK4/6i treatment (**Figure 1E**) and p21 induction is p53-dependent (**Figure 2A**). Using quantitative proteomics of cells arrested in CDK4/6i for different periods of time (see Methods), we observed an increase in p21 protein after two days CDK4/6i treatment, along with increased expression of additional p53 targets, correlating with the increase in cell size (**Figure 2B**).

**Figure 2.**
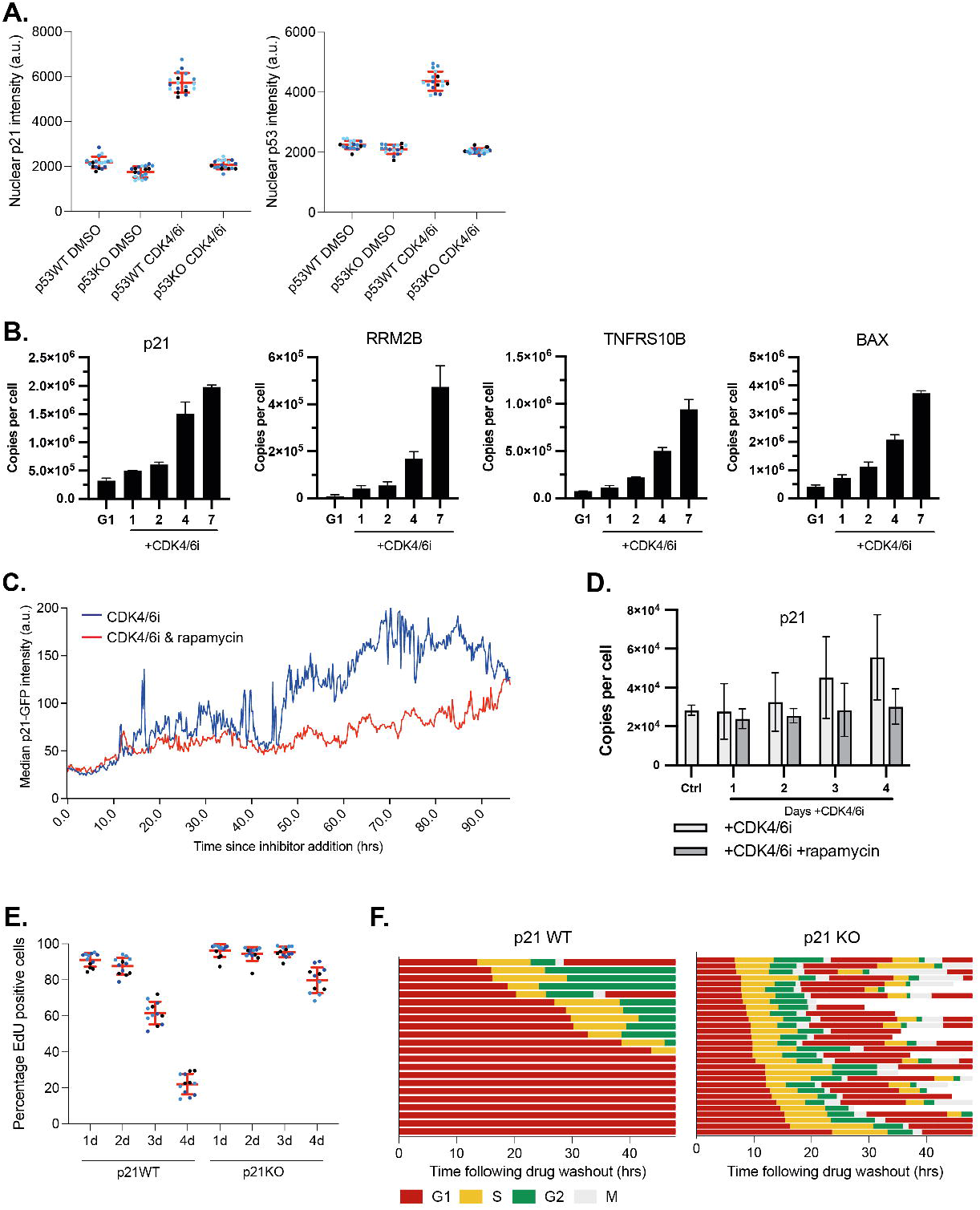
Long-term cell cycle arrest induced by cellular overgrowth is p21-dependent. **A**. Graphs show the increase in p53 and p21 protein during 7d arrest in CDK4/6i. p21 accumulates in a p53-dependent manner. Mean +/- s.d. of n=3 are shown in red. **B**. Bar graphs showing mean protein copies per cell of p53 target proteins in G1 control cells or indicated number of days +CDK4/6i (error bars indicate s.d.). **C**. Graph showing nuclear p21-GFP intensity measured in RPE1 mRuby-PCNA p21-GFP H3.1-iRFP cells after addition of CDK4/6i (blue curve) or CDK4/6i and rapamycin (red curve) to cells. Median p21-GFP intensity of single-cell data are shown: n=78 cells for CDK4/6i and n=66 cells for CDK4/6i and rapamycin (single cell data shown in Supplementary Figure 2B). **D**. Mean copies of p21 protein in control (untreated, proliferating) cells, cells treated with CDK4/6i, and cells treated with CDK4/6i and rapamycin (error bars indicate s.d.). **E**. Graph showing percentage of cells re-entering S-phase (EdU positive) after increasing days of CDK4/6i treatment followed by inhibitor washout in p21 wild-type (WT) and p21 knockout (p21KO) RPE1 cells. RPE1 cells were arrested in 1μM CDK4/6i for the indicated times, inhibitor was then washed out and fresh media containing EdU was added. Cells were fixed and analysed 24h after inhibitor washout. Mean +/- s.d. of n=3 are shown in red. **F**. Graphs show timings of cell cycle phases extracted from time-lapse imaging of p21WT or p21KO RPE1 mRuby-PCNA cells, after release and re-plating from 7d arrest in CDK4/6i or CDK4/6i and rapamycin. Imaging was started 6h after re-plating. Time shown is time from release. Tracks that end prematurely are cells that migrated out of the field of view.

Consistent with these results, imaging of RPE1 mRuby-PCNA p21-GFP cells (Barr et al., 2017) after addition of CDK4/6i or combined CDK4/6i and rapamycin (**Figure 2C**; Supplementary Figure 2A) showed an increase in p21-GFP expression at approximately 48h post-CDK4/6i addition (**Figure 2C**). This increase in p21-GFP expression after 48h of CDK4/6i treatment coincides with an increased fraction of cells remaining in a long-term G0/G1 arrest after two days of CDK4/6i treatment (**Figure 1E**). Rapamycin co-treatment attenuates the increased nuclear p21-GFP fluorescence at 48h. Consistent with this, co-treatment of cells with CDK4/6i and rapamycin prevented the accumulation of p21 protein in total cell extracts (**Figure 2D**) and nuclear p21 in RPE1 (**Supplementary Figure 2B**).

To test whether p21 was required to mediate the long-term G0/G1 arrest after CDK4/6i washout, we treated p21 wild-type (WT) and p21 knockout (KO) cells (Barr et al., 2017) with CDK4/6i for four days and observed that, after CDK4/6i washout, an increased fraction of p21KO cells were able to re-enter S-phase, compared to p21WT cells (**Figure 2E**). Using live-cell imaging of p21WT or p21KO RPE1 mRuby-PCNA cells (Zerjatke et al., 2017), we observed that in the absence of p21, all cells enter S-phase after inhibitor release, have a shorter G1 and cells complete multiple mitoses (**Figure 2F, Supplementary Figure 2C**).

Overall, our results suggest that cellular overgrowth induced by CDK4/6i is sensed by cells and transduced to promote a p21-dependent long-term cell cycle arrest from both G0/G1 and G2.

### Biphasic accumulation of p21 drives long-term cell cycle arrest from G1 or G2

p53-dependent p21 accumulation could explain long-term cell cycle arrest after release from prolonged CDK4/6i treatment, since p21 could block CDK activity and prevent cell cycle re-entry. However, a fraction of cells re-enter S-phase after CDK4/6i washout and withdraw from the cell cycle from G2 (**Figures 2F** (left) and **1H**) in a p21-dependent manner (**Figure 2F**, right). Thus, we investigated how p21 could drive cell cycle exit from G2.

We imaged RPE1 mRuby-PCNA p21-GFP cells after washout of 7-day CDK4/6i treatment to quantify p21-GFP expression after release (**Figure 3A**). In cells that remain arrested in G0/G1, p21-GFP levels remain high (red curves, **Figure 3A, Supplementary Figure 3A**), whereas in cells that re-enter S-phase, p21-GFP is degraded, immediately prior to S-phase entry (Abbas et al., 2008; Nishitani et al., 2008), and is then re-expressed as cells enter G2, leading to a second wave of p21 expression in G2 (blue curves, **Figure 3A, Supplementary Figure 3A-B**). Of note, we did not observe any difference in p21-GFP levels in cells that remain arrested in G0/G1 versus in those that do re-enter S-phase at drug washout (**Supplementary Figure 3A**). In cells co-treated with CDK4/6i and rapamycin, p21-GFP levels are much lower in most cells at the time of release (0h), most cells re-enter S-phase (blue curves, **Figure 3B**) and while p21-GFP does start to accumulate in some cells after S-phase exit, the overall levels are much lower than those observed after treatment with CDK4/6i only and many fewer of these cells arrest in G2 (**Supplementary Figure 3B**; **Figure 1H**).

**Figure 3.**
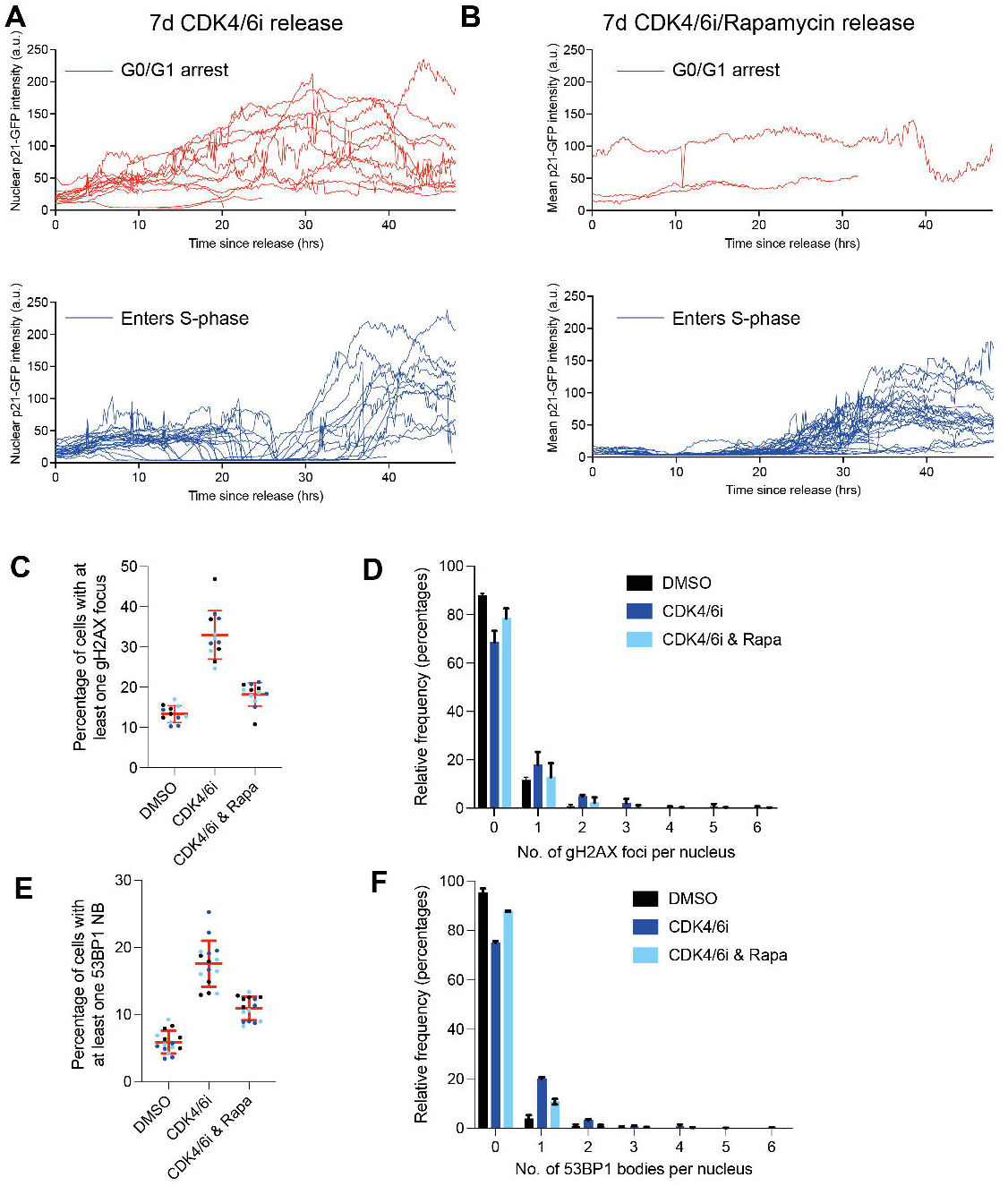
Biphasic accumulation of p21 drives long-term cell cycle arrest. **A, B**. Nuclear p21-GFP intensity measured in RPE1 mRuby-PCNA p21-GFP cells released from 7d CDK4/6i treatment (**A**, n=44 cells) or CDK4/6i and rapamycin treatment (**B**, n=41 cells). Red curves are cells that remain in a G0/G1 arrest, blue curves are cells that enter S-phase. **C-F**. Graphs showing quantification of DNA damage foci 48h after cells are released from a 7d CDK4/6i or CDK4/6i and rapamycin arrest. Left-hand graphs show fraction of nuclei with at least one DNA damage focus measured by large *γ*-H2A.x foci (**C**) or 53BP1 nuclear bodies (**E**). Mean +/- s.d. of n=3 are shown in red. Right-hand graphs show fraction of cells with different numbers of DNA damage foci, measured by large *γ*-H2A.x foci (**D**) or 53BP1 nuclear bodies (**F**). Mean +/- s.d. of n=3 are shown.

Our data suggests that two waves of p21 expression drive long-term cell cycle arrest. We have shown that the first wave of p21 accumulation is induced by cellular overgrowth. Previous work has shown that prolonged arrest in CDK4/6i leads to the downregulation of proteins required for DNA origin licensing and DNA replication (Crozier et al., 2022). This deficiency leads to replication stress in cells that re-enter S-phase after CDK4/6i washout, which could drive p53-dependent p21 expression in G2 after S-phase completion (Barr et al., 2017). If this is the case, then we predict that there should be lower levels of replication stress in cells co-treated with CDK4/6i and rapamycin since p21-GFP does not accumulate to as high a level in these G2 cells. Indeed, co-treatment with rapamycin reduces the amount of DNA damage present in these cells 48h after washout of seven day treatment with CDK4/6i (**Figure 3C-F**). This difference is not due to DNA damage during the G1 arrest, as we have no evidence that DNA damage accumulates during seven day treatment with CDK4/6i (Crozier et al., 2022). Instead, these data suggest that suppressing cellular overgrowth during the prolonged G0/G1 arrest reduces replication stress in cells that re-enter S-phase after inhibitor washout.

These results show that long-term cell cycle arrest is driven by a biphasic p21 response. First, cell overgrowth drives p21 accumulation during the prolonged G0/G1 arrest that prevents cells from re-entering S-phase and second, in cells that re-enter S-phase following CDK4/6i washout, replication stress drives a second wave of p21 expression that promotes cell cycle withdrawal from G2.

### Hyperosmotic media suppresses p21 induction during prolonged G0/G1 arrest

p53-driven p21 expression is activated in response to diverse cellular stresses. However, it is not known what cellular stresses cause the first wave of chronic p21 accumulation induced by cellular overgrowth.

p21 is activated by many stresses (Abbas and Dutta, 2009), including DNA damage (Dulić et al., 1994; El-Deiry et al., 1994), reactive oxygen species (ROS) (Barnouin et al., 2002) and osmotic stress (Kishi et al., 2001). Markers of DNA damage, including *γ*-H2A.x foci (Crozier et al., 2022; Wang et al., 2022) and 53BP1 nuclear bodies, do not increase during prolonged CDK4/6i-mediated G0/G1 arrest (Crozier et al., 2022; Wang et al., 2022) (**Supplementary Figure 4A**), making nuclear genome damage an unlikely culprit for p21 induction. Increasing antioxidant capacity by treatment with the ROS scavenging small molecule, N-acetyl cysteine (NAC), failed to prevent p21 induction (**Supplementary Figure 4B**). ROS is an umbrella term for multiple chemical species, including superoxide and hydrogen peroxide, which have differing reactivities towards NAC. The NRF2 transcription factor is a master regulator of the cellular antioxidant response (Tonelli et al., 2017), upregulating many ROS scavenging and repair pathways, including glutathione biosynthesis and the thioredoxin antioxidant system. NRF2 activity can be increased using the small molecule TBE-31 (Dinkova-Kostova et al., 2010), which stabilizes NRF2 by inhibiting its E3 ligase KEAP1 (Dinkova-Kostova et al., 2002). Treatment of RPE1 cells with TBE-31, leads to increased expression of the NRF2 target gene NQO1 (**Supplementary Figure 4C**) demonstrating NRF2 activation, but fails to prevent p21 induction during CDK4/6i treatment (**Supplementary Figure 4C-D**). Therefore, our data suggest that, although CDK4/6i has been shown to increase ROS^15,33,34^, in RPE1 cells increased ROS levels do not appear to be critical for p21 induction.

To investigate if osmotic stress could contribute to p21 induction, we tested whether extracellular osmolyte concentration can modulate p21 levels in CDK4/6i treated cells. Cells arrested for one day in CDK4/6i were cultured for an additional three days in CDK4/6i with media with different osmolarities: hypertonic (703 mOsm/kg), hypotonic (179 mOsm/kg) and isotonic (300 mOsm/kg). Critically, nutrient concentrations were identical between media. The only varying component was the concentration of D-sorbitol, either absent (hypotonic), restored to basal levels (isotonic), or present in excess (hypertonic). Addition of hypertonic media to asynchronous cells increases p21 levels, as expected (Kishi et al., 2001) (**Supplementary Figure 4E**). However, in CDK4/6i treated cells, hypertonic media suppresses p21 induction, measured by both flow cytometry (**Figure 4A**) and microscopy (**Figure 4B**). Cell size was largely unaffected (**Supplementary Figure 4F**), presumably because of the timescale of the treatment with hypertonic media (three days) and homeostatic regulatory volume increase (Hoffmann et al., 2009).

**Figure 4.**
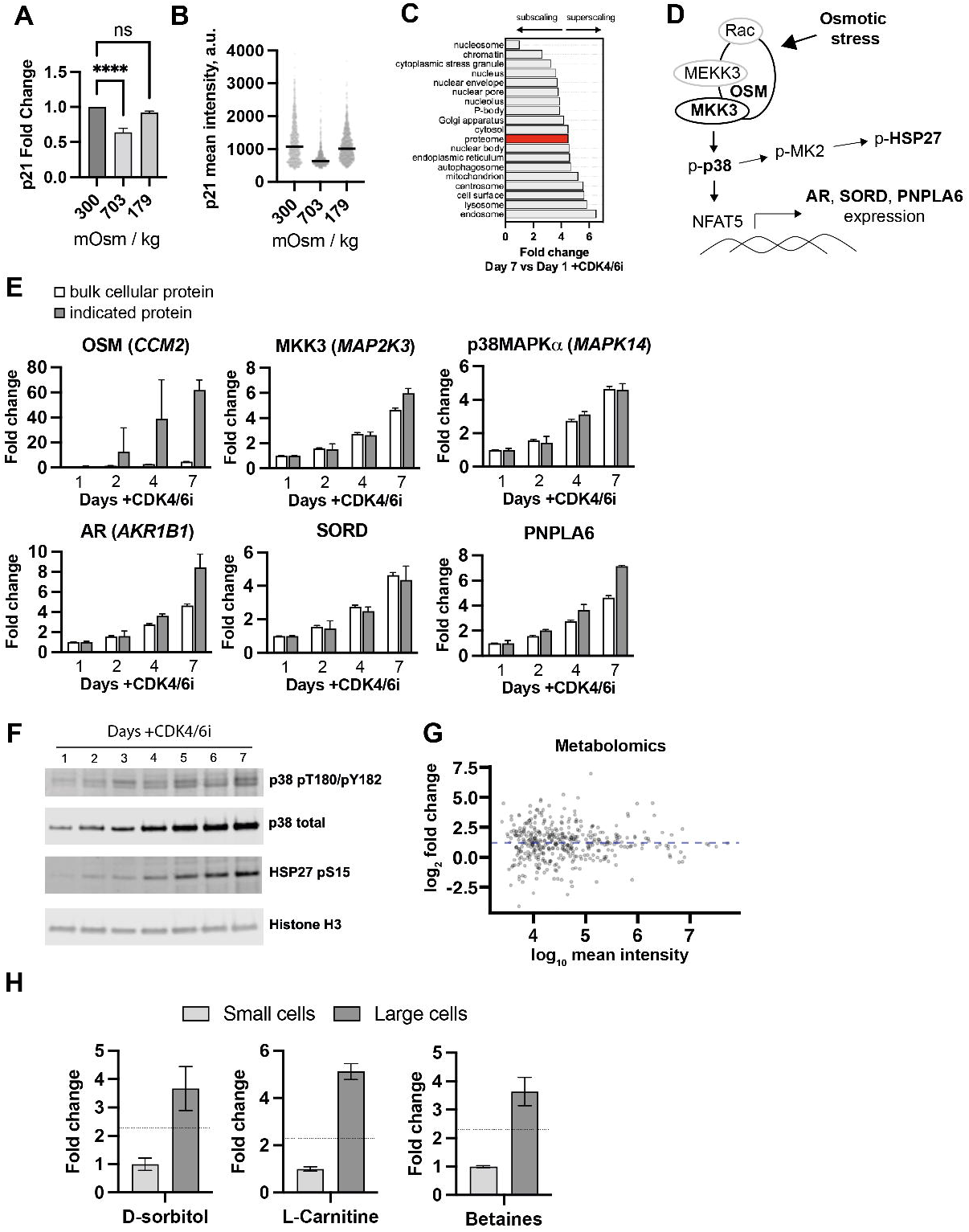
Cell growth during prolonged G1 arrest activates an osmoadaptive response. **A**. Flow cytometry analysis of RPE1 cells treated with CDK4/6i for 4 days and either isotonic (300 mOsm/kg), hypertonic (703 mOsm/kg) or hypotonic media (179 mOsm/kg). Cells were immunostained for p21. **B**. Quantitative microscopy analysis of RPE1 cells treated as in (A) and immunostained for p21. **C**. Median fold change comparing day 7 versus day 1 CDK4/6i for proteins localized to indicated subcellular compartments. “Proteome” indicates fold change for average protein. **D**. An abbreviated schematic of the osmotic shock and osmoadaptation pathway. **E**. Comparison of protein copy fold change of indicated proteins and bulk cellular protein across indicated days of CDK4/6i treatment. Error bars show s.e.m. (n = 2-3). **F**. Immunoblot analysis of RPE1 cells treated with CDK4/6i for the indicated days. **G**. Fold changes for metabolites (n = 416) comparing day 4 versus day 1. X-axis shows mean intensity across conditions/replicates. **H**. Fold change of osmolytes, D-sorbitol, L-carnitine, and betaines. These osmolytes were detected with confidence levels of 2, 1, and 1, respectively, as defined by the Metabolomics Standards Initiative (MSI).

These data show that altering the osmolarity of extracellular media can limit the induction of p21 expression. Next, we wanted to further investigate the possibility that cell overgrowth may induce, or exacerbate, osmotic stress.

### Cell growth during prolonged G1 arrest activates an osmoadaptive response

Osmotic stress occurs when extracellular and intracellular osmolarities differ. In CDK4/6i-treated cells, a change in the osmolarity of the culture media is unlikely. However, it is possible that cell overgrowth challenges osmotic balance by affecting intracellular osmolarity. For example, intracellular osmotic forces play key roles in morphological changes in cells cultured in isotonic media, including cell flattening during cell-matrix attachment^51^ and cell rounding during mitosis^52^. Proteomic analysis (**Supplementary Table 1**) demonstrated that excessive growth is accompanied by widespread proteome remodeling characterized by imbalanced scaling of subcellular organelles (**Figure 4C**) (Lanz et al., 2022), which may also threaten osmotic balance. Cells respond to hyperosmotic stress by regulatory volume increase (Hoffmann et al., 2009). Signal transduction and gene expression changes lead to accumulation of intracellular osmolytes (Burg and Ferraris, 2008), which buffer cells against extracellular hyperosmotic conditions. If cells experience osmotic stress in becoming aberrantly large, then we should observe induction of osmotic adaptation pathways.

Osmotic stress triggers a signaling pathway (**Figure 4D**) that includes activation of the key stress-associated p38 mitogen-activated protein kinase (Uhlik et al., 2003). Many proteins in this pathway were detected in our proteomics dataset (**Figure 4D**, in bold). The scaffolding protein OSM (*CCM2*) recruits Rac, MEKK3 and MKK3 to activate p38MAPK. OSM is present at ∼3,000 copies in day one CDK4/6i cells and increases to ∼180,000 copies at day seven CDK4/6i (**Figure 4E**). OSM is thus a superscaler (Lanz et al., 2022), increasing disproportionately higher (∼60-fold) than increases in bulk protein content (∼4.5-fold, white bars in **Figure 4E**). The p38MAPK activating kinase, MKK3 (MAP2K3), also superscales. The levels of total p38MAPK increase (**Figurs 4D-E**), scaling with size (**Figure 4E**). Therefore, our proteomics data suggests that the osmotic stress pathway is activated in overgrown CDK4/6i treated cells. To explore this further, we assayed active p38MAPK, by measuring phospho-p38MAPK pT180/pY182 and found that p38MAPK activity also increases with size (**Figure 4F**). Furthermore, phospho-HSP27 pS15, a downstream substrate of p38MAPK (Cuenda and Rousseau, 2007), increases with size (**Figure 4F**), consistent with activation of the p38MAPK pathway.

p38MAPK activates the transcription factor NFAT5 (Ko et al., 2002) to promote the expression of numerous genes important in osmotic regulation (Burg and Ferraris, 2008), including aldose reductase (AR), sorbitol dehydrogenase (SORD) and the phospholipase - neuropathy target esterase (PNPLA6), which all scale or superscale with cell size (**Figure 4E**). AR produces sorbitol from glucose and SORD converts sorbitol to fructose, thereby controlling intracellular levels of D-sorbitol, which is critical for osmotic regulation in mammalian cells (Burg and Ferraris, 2008). To measure levels of D-sorbitol and other metabolites important for intracellular osmolarity regulation, we performed untargeted metabolomics comparing day four (large) and day one (small) CDK4/6i-treated cells. 416 metabolites were putatively assigned based on accurate mass. Equal numbers of cells from each condition were analyzed. Consistent with size-scaling of the metabolome, metabolites increase 2.3-fold on average, which is proportional to the increase in volume and protein content (**Figure 1B-C**, ∼2.5-fold). However, individual metabolites vary in scaling behaviour (**Figure 4G-H**). Levels of D-sorbitol increase 3.7-fold (**Figure 4H**), i.e., more than average metabolites increase (**Figure 4H**, indicated as dotted lines), suggesting that the changes in AR and SORD (**Figure 4E**), enzymes crucial for sorbitol anabolism and catabolism, produce a net increase in the intracellular concentration of sorbitol. Other osmolytes also superscale, including L-carnitine and betaine (**Figure 4H**). The osmolyte myo-inositol is isobaric with other hexoses and therefore its scaling behaviour cannot be accurately determined using this assay.

Based on these observations, we conclude that while the metabolome scales with size, key intracellular osmolytes superscale, consistent with an active osmoadaptive response in large cells.

### p38MAPK activity is required for p21 induction during G1 arrest and cell cycle defects following release from G1

p38MAPK is a core signaling node downstream of osmotic and other cellular stresses (Cuenda and Rousseau, 2007) to activate p53 transcriptional activity and increase p21 levels (Kishi et al., 2001). To test if p38MAPK activity plays a critical role in the first wave of p21 induction, cells were treated with either CDK4/6i alone, or CDK4/6i and a small molecule inhibitor of p38MAPK (SB-203580, p38i)(Cuenda et al., 1995). Whilst CDK4/6i + p38i treated cells grew to a similar size to CDK4/6i treatment alone (**Figure 5A, Supplementary Figure 5A**), p21 induction following CDK4/6i is almost completely prevented by the co-treatment with p38i (**Figure 5B, Supplementary Figure 5B**). Osmotic stress can also activate related stress activated JNK kinases (Galcheva-Gargova et al., 1994). However, co-treatment with a small molecule JNK inhibitor (JNK-IN-8), which targets JNK1/2/3, failed to prevent p21 induction (**Supplementary Figure 5C**).

**Figure 5.**
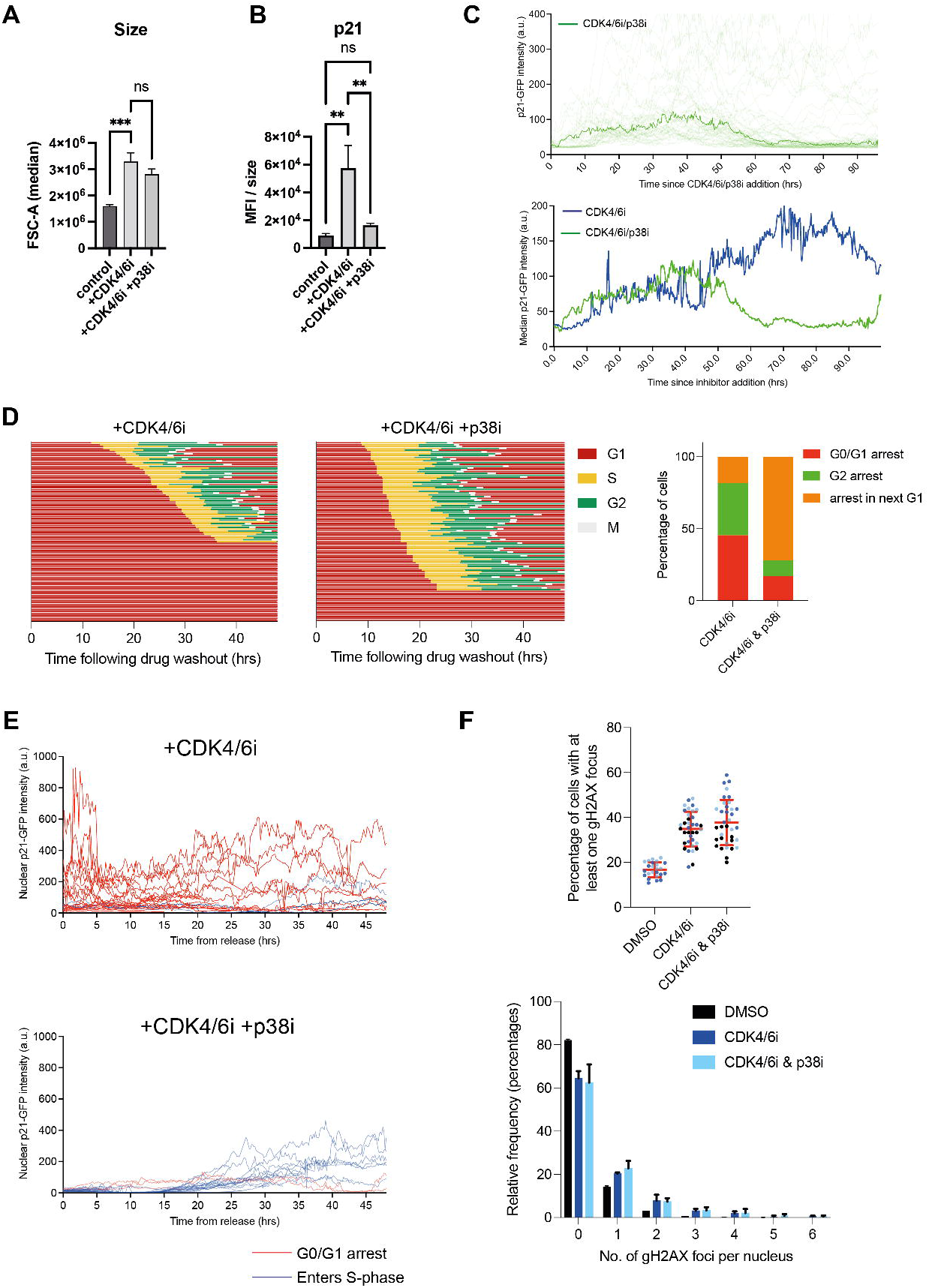
p38MAPK activity is required for p21 induction during G0/G1 arrest and cell cycle defects following release from G1. **A**. Forward scatter of cells treated with CDK4/6i or with co-treatment with p38i. **B**. Cells were treated as in (A), immunostained for p21, and assayed by flow cytometry. MFI = geometric mean of fluorescence intensity. MFIs were normalized to size (forward scatter). **C**. Graph showing nuclear p21-GFP intensity measured in RPE1 mRuby-PCNA p21-GFP H3.1-iRFP cells after addition of CDK4/6i and p38i (SB203580; green curve) or CDK4/6i alone (blue curve) to cells. Median p21-GFP intensity of single-cell data are shown: n=87 cells for CDK4/6i/p38i (single cell data shown in faint green curves in top graph). **D**. Graphs show timings of cell cycle phases extracted from time-lapse imaging of RPE1 mRuby-PCNA p21-GFP cells after release from 7d arrest in CDK4/6i or CDK4/6i and p38i (30 μM SB-203580). In these experiments, cells were released into fresh media with no STLC and so were able to complete mitosis and enter the next G1. Tracks that end prematurely are cells that migrated out of the field of view. Right-hand graph shows quantification of cell cycle fates after release. **E**. Nuclear p21-GFP intensity measured in RPE1 mRuby-PCNA p21-GFP H3.1-iRFP cells released from 7d CDK4/6i treatment (n=24 cells) or CDK4/6i and p38i treatment (n=22 cells). Red curves are cells that remain in a G0/G1 arrest, blue curves are cells that enter S-phase. **F**. Graphs showing quantification of DNA damage foci 48h after cells are released from a 7d CDK4/6i or CDK4/6i and p38i (SB-203580) arrest. Top superplot shows fraction of nuclei with at least one gammaH2A.X large focus. Bottom graph shows fraction of cells with different numbers of large gammaH2A.X foci. Mean +/- s.d. are shown.

To observe when during a CDK4/6i-induced G0/G1 arrest p38i acts to suppress p21, we used live-cell imaging of RPE1 mRuby-PCNA p21-GFP cells treated with CDK4/6i alone or CDK4/6i + p38i (**Figure 5C**). As shown before (**Figure 2C**), average p21-GFP levels increase at ∼48h of CDK4/6i treatment. By contrast, this induction of p21-GFP is not only suppressed in CDK4/6i + p38i treated cells but actually decreases around the same timepoints (**Figure 5C**). We conclude that p38MAPK activity is required for p21 induction during a prolonged G0/G1 arrest.

Next, we investigated if suppression of p21 induction by p38i leads to rescue of long-term cell cycle arrest. RPE1 mRuby-PCNA p21-GFP cells were treated for seven days with either CDK4/6i alone, or with CDK4/6i + p38i. As shown in **Figure 5D**, CDK4/6i-treated cells released from G1 arrest were delayed in S-phase entry (∼55%) or did not re-enter the cell cycle (∼45%). Of those that entered S-phase, ∼29% did not enter mitosis, arresting in G2 (Figure 5D). Co-treatment with CDK4/6i and p38i reduced both S-phase entry delay and the G2 arrest (**Figure 5D**). This rescue of G2 arrest was reproduced in the RPE FUCCI cell line and with three structurally distinct inhibitors of p38MAPK: SB-203580 (p38i), VX-745, BIRB-796 (**Supplementary Figure 5D**).

We attribute the reduction in p21 induction during G0/G1 arrest as the reason why more cells co-treated with CDK4/6i and p38i re-enter S-phase and why these cells have a shorter G1 (**Supplementary Figure 5E**). However, many of these cells still arrest in the next G1 (**Figure 5D**). To understand why CDK4/6i and p38i co-treatment is not able to fully rescue the long-term cell cycle arrest phenotypes, we used live cell imaging to measure p21-GFP expression in RPE1 mRuby-PCNA p21-GFP after release from seven day CDK4/6i or CDK4/6i and p38i treatment. As described, p21-GFP levels are lower after washout of CDK4/6i + p38i and the majority of these cells re-enter S-phase (blue curves, **Figure 5E**). However, after completing S-phase, cells released from dual CDK4/6 and p38 inhibition upregulate p21-GFP during G2 (**Supplementary Figure 5F**). We observed this increase in p21 protein after inhibitor removal also in fixed cells (**Supplementary Figure 5G**). Based on our previous observations (**Figure 3**) we suspected that this is because CDK4/6i and p38i co-treated cells remain overgrown and so remain sensitive to increased replication stress during the first S-phase after release from CDK4/6i. We quantified the level of DNA damage in cells released from CDK4/6i + p38i treated cells and saw that the levels of DNA damage were equivalent to those in cells treated with CDK4/6i alone (**Figure 5F**). This suggests that cellular overgrowth may be contributing to problems during mitosis, which may contribute to arrest in the next G1.

Our data show that p38MAPK inhibition can prevent the first wave of p21 expression - during a prolonged CDK4/6i-induced G0/G1 arrest. However, since cells are still aberrantly large, the second wave of p21 expression, that is dependent on replication stress, persists, and therefore cells treated with combined CDK4/6i and p38i still withdraw from the cell cycle in G2 or the subsequent G1.

## DISCUSSION

Cell growth and proliferation are coupled to regulate cell size. Uncoupling these processes has major, negative consequences on cells (Cheng et al., 2021; Lanz et al., 2022; Lengefeld et al., 2022; Neurohr et al., 2019) but how this is controlled at a molecular level remains under investigation. We have shown that during prolonged CDK4/6 inhibition, cells grow abnormally large. This cellular overgrowth promotes long-term cell cycle arrest, even after CDK4/6i removal. This long-term cell cycle arrest is p53-p21 dependent and is driven by, at least, two events that promote distinct waves of p53-dependent p21 protein expression (**Figure 6**). The first p21 wave occurs during CDK4/6i treatment - where osmotic stress can activate p38MAPK, which promotes downstream p21 expression. The second p21 wave occurs after removal of CDK4/6i in a fraction of cells that re-enter S-phase. In these cells, aberrant size generates replication stress which promotes additional p21 expression after completing the first S-phase, causing cells to withdraw from the cell cycle in G2, or the next G1. The levels of p21 therefore integrates signals over multiple days from chronic osmotic stress and acute replication stress to determine cell fate.

**Figure 6.**
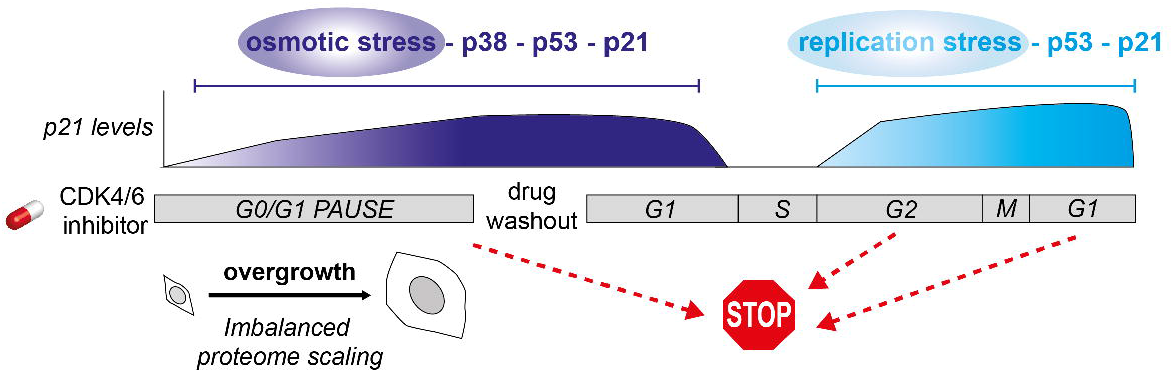
Model for long-term inhibition of proliferation after CDK4/6i removal. Prolonged treatment of cells with CDK4/6i promotes cellular overgrowth. Cell overgrowth is accompanied by an osmotic stress response in cells that induces a p38MAPK-dependent, p53-dependent p21 expression that promotes permanent cell cycle arrest after CDK4/6i removal. Overgrown cells that re-enter S-phase after CDK4/6i removal experience replication stress which promotes further p21 expression resulting in permanent arrest in G2, or the subsequent G1.

The cytoplasmic dilution model (Neurohr et al., 2019) proposes that aberrantly large cells face a fitness penalty due to inefficiencies in gene expression and biochemical reactions in a dilute cytoplasm. A reduction in large macromolecular crowding in an expanding cytoplasmic volume must be accompanied by an increase in more freely diffusing solutes to maintain osmotic balance. We demonstrated that large cells showed activation of an osmolyte biosynthesis pathway via p38, leading to superscaling of osmolytes in cells. Here p38 activation serves two roles: regulation of intracellular osmolyte concentration via changes in gene expression and stress signal transduction via p53.

Cell growth during an unperturbed G1, and during a CDK4/6i-induced G1 arrest, is known to cause allometric scaling of subcellular organelles, including subscaling of chromatin proteins (especially histones) and superscaling of lysosomes (Lanz et al., 2022). We observe the same trend in our proteomics data (Figure 4C). While osmotic stress and the cytoplasmic dilution model are consistent, studies thus far have not demonstrated why either phenomena occur in large cells. We favour the possibility that widespread proteome remodeling and loss of proteostasis is one source of osmotic stress. Indeed, unequal gene dosage perturbs protein complex subunit expression and stoichiometry, leading to hypo-osmotic stress (Tsai et al., 2019). Subunit stoichiometry uncoupling has also been implicated in cellular aging in yeast (Janssens et al., 2015). However, this model will require robust evaluation and testing.

Like G2 progression, cell cycle re-entry following release from prolonged CDK4/6i treatment is also heterogeneous, with some cells withdrawing from the cell cycle in G0/G1 and others able to re-enter S-phase and withdraw after S-phase completion. However, in these experiments, we were unable to establish the cause of this heterogeneity. We observed no significant difference between p21-GFP levels or nuclear area in cells that re-entered S-phase versus those that remained in G0/G1. Other mechanisms might play more important roles here. For example, the ratio between CyclinD1 and p21 is important in determining whether cells proliferate or not (Yang et al., 2017). Cyclin D1 superscales with size during CDK4/6i treatment (**Supplementary Figure 6A**) and so it is possible that cells that re-enter S-phase may have a higher Cyclin D1/p21 ratio.

The lack of apparent DNA damage from cells withdrawing from G0/G1 is crucial (Crozier et al., 2022; Manohar et al., 2022; Wang et al., 2022) (**Supplementary Figure 4A**). Cells that enter S-phase and undergo replication stress will trigger a DNA damage response, which was demonstrated to activate NF-kappa-B and induce a secretory phenotype that promotes proliferation and migration of tumour cells (Chien et al., 2011; Rodier et al., 2009). In contrast, a p53-associated secretory phenotype lacking DNA damage induces potent immunosurveillance while lacking tumour promoting characteristics (Wang et al., 2022). Interestingly, chronic p38 activity is sufficient to promote a senescence associated secretory phenotype, which can operate independently from DNA damage signaling (Freund et al., 2011). Thus, by activating p38, cell overgrowth could promote two hallmark characteristics of senescence: permanent cell cycle arrest via p53 and a secretory phenotype. The secretory phenotypes of G0/G1 versus G2 arrested cells, and the role of p38, will be important to determine

CDK4/6 and p38MAPK have previously been implicated in the regulation of cell size (Tan et al., 2021). There, partial inhibition of CDK4/6 activity during G1 leads to a larger target size for the cell by increasing both the duration and the rate of growth. p38MAPK then acts downstream of changes in cell size whereby p38 activity increases in cells below their target size and prevents cell cycle progression. The sensor(s) that detect changes in cell size and lead to p38 activation remain unknown. In our experiments, inhibition of CDK4/6 activity leads to complete arrest in G0/G1, and we attribute continued cell growth as a secondary effect of blocking proliferation and not a direct effect of CDK4/6i. This was confirmed by arresting cells in CDK4/6i for four days, washing out the inhibitor and leaving for three days and observing that the cells that remain arrested in G0/G1 grow to the same size as those kept in CDK4/6i for the full seven days (**Supplementary Figure 6B**). p38 is thus activated downstream of cell overgrowth and osmotic stress. It is possible that partial inhibition of CDK4/6 activity in proliferating G1 cells by Tan et al. also induces a level of osmotic stress and thereby activates p38.

An assumption here that cell overgrowth is a secondary effect of inhibiting cell proliferation, and not a direct consequence of CDK4/6i, is that any inhibitors that block cell cycle entry would also lead to aberrant cell overgrowth. Indeed, recent work has shown that inhibition of CDK7 with ICEC0942 also arrests cells in G0/G1 and leads to cell overgrowth that contributes to long-term cell cycle arrest (Wilson et al., 2021). However, one can argue that CDK7, being part of the CDK-activating kinase (CAK), is upstream of CDK4/6 and therefore is acting through the same pathway. Therefore, it remains to be seen if drugs that act to block cell proliferation, without inducing DNA damage, could also induce long-term cell cycle arrest due to cell overgrowth (see accompanying paper by Foy et al., 2022).

It is clear from our data that the increase in replication stress observed in the first S-phase after removal of CDK4/6 inhibition (Crozier et al., 2022) is, at least in part, due to cells being aberrantly large. Increased replication stress after CDK4/6i removal has been attributed to decreased protein levels of DNA replication licensing factors and a subsequent decrease in replication origin licensing. We observe a similar increase in nuclear area as observed for cell volume during CDK4/6i, although we were unable to measure nuclear volume in a high-throughput way as when we removed the cytoplasm, we found that nuclei in CDK4/6i treated cells were extremely fragile and prone to lysing. However, we hypothesise that increased nuclear area, together with subscaling of DNA replication licensing factors (**Supplementary Table 1**) (Crozier et al., 2022), could cause an overall decreased concentration of nuclear proteins leading to lower efficiency in DNA origin licensing. In addition, in recent work, it has been shown that cellular overgrowth makes cells more sensitive to DNA damage during the G0/G1 arrest (Manohar et al., 2022). This was, at least in part, attributed to a reduced capacity for p53 to respond to DNA damage occurring in these arrested cells. Increased sensitivity to genome damage (Manohar et al., 2022) plus a reduction in DNA origin licensing (Crozier et al., 2022) would both contribute to increased replication stress during the first S-phase after release. There may also be changes in chromatin organisation during CDK4/6i treatment and/or cell overgrowth that could contribute to increased replication stress and this remains to be fully explored.

Whilst our mechanistic investigation here has been in non-transformed cells, to remove the unknown effects of confounding mutations/amplifications/deletions in key signalling pathways, our data has consequences for how we should use CDK4/6i in the clinic. For example, 60% of CDK4/6i-resistant breast cancer patients have mutations in TP53 (Wander et al., 2020). The authors found that p53 knockout cells still arrested in CDK4/6i and so ruled p53 out as contributing to therapy resistance. We have also observed that p21 is not required for cells to arrest in G0/G1 with CDK4/6i (Pennycook and Barr, 2022). However, the data presented here and that of others (Crozier et al., 2022; Wang et al., 2022) show that a functional p53-p21 pathway is required after CDK4/6i removal to mediate long-term cell cycle arrest. Mutations in the p38-osmotic stress pathway were not observed in the Wander et al. study but would not be required to mediate resistance if downstream TP53 is mutated. Moreover, redundancy between p38 isoforms (alpha, beta, gamma, delta) may mean the likelihood of genetic lesions that render loss of p38 function is less likely as compared with mutation of a single tumour suppressor gene such as p53.

Finally, p53 loss is predicted to confer more resistance than p38 loss because it also antagonizes the replication-stress induced DNA damage response. Because the secretory phenotype of senescent cells varies depending on DNA damage (Wang et al., 2022), the lack of DNA damage in p38-dependent long-term cell cycle arrest suggests that enhancing p38 activity with CDK4/6 inhibition could result in a clinically beneficial secretory phenotype by promoting immunosurveillance without pro-proliferative and pro-migratory paracrine signaling to neighbouring tumour cells. It will now be important to test these ideas in vitro and in animal models.

## MATERIALS AND METHODS

### Cell culture

hTERT-RPE1 mRuby-PCNA cells were a generous gift from Jörg Mansfeld (Institute of Cancer Research, London) and their generation is described in (Zerjatke et al., 2017). hTert-RPE1 mRuby-PCNA p21-GFP and hTert-RPE1 mRuby-PCNA p21KO1A were previously described in (Barr et al., 2017). hTert-RPE1 mRuby-PCNA p21-GFP H3.1-iRFP cells were generated using AAV-mediated recombination to label endogenous Histone H3.1 (locus *HISTH31E*) at the C-terminus, as described in. hTERT-RPE1-FUCCI cells were published previously (Krenning et al., 2014). All cell lines were maintained in DMEM (41966; Gibco) supplemented with 10% heat-inactivated FBS and 1% penicillin-streptomycin (both from Sigma) at 37°C and 5% CO2. Cells were screened for mycoplasma by PCR every 1–2 months. hTERT-RPE1 cells (ATCC) were maintained in 1:1 DMEM/F-12 (10565018, Thermo) supplemented with 9% FBS (10270106, Thermo).

### Inhibitors and reagents

The following inhibitors were used in this study: Palbociclib (PD-0332991, hydrochloride salt, MedChemEx-press, HY-50767A, Merck PZ0383); Rapamycin (MedChemExpress, Cat. AY-22989), JNK-I-8 (Merck SML1246), SB-203580 (MedKoo, 574872, Merck 559389), VX-745 (Tocris 3915), BIRB-796 (Tocris 5989), and S-Trityl-L-cysteine (STLC; Sigma Aldrich, #164739). TBE-31 was a gift from Albena Dinkova-Kostova (University of Dundee). Inhibitor stocks were diluted in DMSO, aliquoted and stored at −20’C. N-acetyl cysteine (A7250, Merck) was prepared in water.

### Fixed-cell experiments to analyse release from CDK4/6 inhibitors

Cells were plated on black 384-well CellCarrierUltra (PerkinElmer) plates one day before adding inhibitors at a density of 2.5×10^4^ cells/ml in 20 μl of media/well. After cells had been incubated in inhibitors for the required number of days, depending on the experiment, inhibitor containing media was aspirated and cells washed four times in PBS, using an automated liquid handler – 50TS microplate washer (Biotek). Fresh media was then added to cells containing 10 μM EdU, in order to label cells as they enter S-phase. After 24 or 48h further incubation, cells were fixed and immunostained, as described below.

### Fixed-cell experiments to quantify DNA damage after release

Cells were plated at a concentration of 1×10^4^ cells/ml in 6 well plates. After 6hr, when cells had attached, inhibitors were added to cells for seven days, with media and inhibitors being refreshed after 3 days. After seven days, inhibitors were washed from cells by washing four times in PBS, cells were trypsinised, resuspended in fresh media, centrifuged at 1,000xg for 3 mins, media aspirated, cells resuspended in 2ml of fresh media and 40ul of media was plated per well of a black 384-well CellCarrierUltra (PerkinElmer). Cells were left for 48h to release, then fixed and immunostained, as described below.

### Fixed-cell experiments to quantify protein levels after inhibitor treatment

Cells were plated on black 384-well CellCarrierUltra (PerkinElmer) plates one day before adding inhibitors at a density of 1.25×10^4^ cells/ml in 20 μl of media/well. After cells had been incubated in inhibitors for the required number of days, cells were fixed and immunostained, as described below.

### Immunofluorescence

For immunostaining in 384 well plates, cells were fixed by adding and equal volume of 8% formaldehyde in PBS to media in wells to give a final concentration of 4% formaldehyde. Cells were fixed for 15 minutes at RT and washed three times with PBS. Permeabilization in PBS 0.5% Triton X-100 for 15 min was followed by blocking in 2% BSA in PBS (blocking buffer) for 1 h. Cells were incubated with primary antibodies diluted in blocking buffer at 4°C overnight and washed three times with PBS, then incubated with a 1:1000 dilution of Alexa-labelled secondary antibodies (ThermoFisher), also diluted in blocking buffer for 1h at RT. Finally, cells were incubated with Hoechst 33258 (1ug/ml) for 15 min at RT, followed by 3 washes in PBS. All washes and aspirations were performed on the automated liquid handler (50TS microplate washer (Biotek)). Primary antibodies used for immunostaining in this study were: p21 (BD 556430, 1:500), p21 (Invitrogen MA5-14949: 1:1000), p53 (CST 2527, 1:500), *γ*-H2A.x (CST 2577, 1:2000), 53BP1 (CST 4937,1:1000).

For EdU staining, a final concentration of 10 μM EdU was added to growth media. After fixing, permeabilising and blocking (as detailed above), the Click-iT reaction was performed as follows: cells were incubated in the dark, for 30 min at RT in a solution of 100 mM Tris-HCl pH 7.5, 4 mM CuSO4, 5 μM sulfo-cyanine-3-azide and 100 mM sodium ascorbate. Cells were then washed three times in PBS, and either immunostained with primary and secondary antibodies (detailed above) or directly incubated for 15 min at RT with 1 μg/ml Hoechst 33258, and then washed three times in PBS. Fixed cell imaging of 384 well plates was performed on the Operetta CLS (PerkinElmer) high-content microscope using the 20x N.A. 0.8 objective.

### Image quantification

All fixed cell image analysis was performed in Harmony automated image analysis software (PerkinElmer). Data were plotted in GraphPad Prism 7 and repeat experiments plotted as Superplots(Lord et al., 2020), where appropriate.

To calculate nuclear intensities and cell and nuclear area, nuclei were identified and segmented based on Hoechst staining. Poorly segmented, nuclei at the edge of the field of view, mitotic and dead cells were excluded from analysis based on a combination of features measuring area, roundness and Hoechst intensity. Nuclear intensities and nuclear area were then quantified in well-segmented nuclei.

To calculate the percentage of EdU positive cells, EdU intensity was quantified in well-segmented nuclei. EdU positive cells were identified via setting a threshold based on the biphasic population.

To identify DNA damage foci with *γ*-H2A.x or 53BP1 immunostaining, the spot finding function B of the Harmony software was used. Small foci labelled by these antibodies that are frequently visible in S-phase cells were excluded based on their smaller size and lower staining intensity.

### Live imaging

#### CDK4/6i release experiments

hTert-RPE1 Ruby-PCNA or hTert-RPE1 Ruby-PCNA p21-GFP H3.1-iRFP cells were seeded at a density of 2.5×10^4^ cells/ml in 20 μl of media/well in 384-well CellCarrier Ultra plates (PerkinElmer) or 8-well Ibidi chambers one day before adding inhibitors. After seven day incubation in inhibitors, wells were washed four times in PBS and inhibitor-containing media was replaced with fresh phenol-red free DMEM with 10% FBS and 1% P/S before live imaging was started. Live imaging was performed on either the Operetta CLS high-content microscope (PerkinElmer) at 20x N.A. 0.8 in confocal mode or the Olympus ScanR confocal microscope at 20x N.A. 0.7. In both cases, cells were imaged under climate-controlled conditions at 37°C and 5% CO_2_. Cells were acquired every 10 or 12 mins. To quantify the timing of release and subsequent cell cycle phase lengths, experiments were analysed manually using FIJI. To quantify p21-GFP levels after release, images were analysed using Nuclitrack(Cooper et al., 2017).

#### p21-GFP quantification experiments

To quantify p21-GFP levels after addition of inhibitors to asynchronous cells, cells were plated in phenol-red free DMEM (10% FBS, 1% P/S) at a density of 2.5×10^4^ cells/ml in 20 μl of media/well, one day prior to the start of the experiment. The next day, inhibitors were added and media made up to a final volume of 100ul per well. Imaging was started immediately on the Operetta CLS microscope in confocal mode (PerkinElmer), using 20x N.A. 0.8 and acquiring images every 10-12 mins. Cells were maintained at 37°C and 5% CO_2_ throughout imaging. To quantify p21-GFP levels after inhibitor addition, images were analysed using Nuclitrack(Cooper et al., 2017).

#### Immunofluorescence flow cytometry

Cells were fixed in 2% formaldehyde for 15 min at 37 °C, washed in flow buffer (1% BSA in DPBS), and resuspended in ice-cold 90% methanol. For immunostaining, methanol-treated cells were washed with flow buffer, resuspended in immunostain solution containing 1:50 anti-p21 AlexaFluor 647 conjugate (Cell Signaling Technologies, #8587) in flow buffer and incubated at RT for 30 min. Cells were then washed with flow buffer and assayed using a Novocyte flow cytometer (ACEA Biosciences).

#### Modulation of extracellular osmolarity

DMEM/F12 was diluted 1:1 with sterile water and supplemented with sodium bicarbonate to 29 mM final (Thermo 25080094). Hypo-osmotic (179 mOsm/kg), isotonic (300 mOsm/kg) and hyper-osmotic (703 mOsm/kg) media was prepared by adding varying amounts of sorbitol (Sigma S1876). Flow cytometry analysis was performed as described above. Immunofluorescence microscopy was performed by culturing RPE1 cells on N1.5 13 mm coverslips (VWR), fixing cells with 4% formaldehyde (Thermo) at 37 °C for 15 min, permeabilizing with 0.1% Triton for 5 min, blocking with 1% BSA in TBS, incubating with primary antibody (1:200 dilution in 1% BSA in TBS) for 16 hr at 4°C, and then incubating with secondary antibodies (1:200 dilution in 1% BSA in TBS). Cells were then stained with DAPI, mounted on glass slides using ProLong Glass (Thermo) and imaged using a DeltaVision microscope (Imsol). Antibodies used were anti-p21 (CST #2947), anti-tubulin (DM1A, Merck), goat anti-rabbit AlexaFluor 555 (abcam, ab150078) and donkey anti-mouse AlexaFluor 488 (Thermo, A32766).

Cell volumes were measured using a HoloMonitor M4 in either Sarstedt Lumox, or Ibidi μ-plate glass-bottomed 24 well plates and the associated hololids. Optical volumes were determined using the holomonitor app suite.

#### Proteomics

Cells were scraped in 2% SDS lysis buffer containing phosphatase inhibitors (PhosStop, Roche) and protease inhibitors (Complete EDTA-free, Roche). Extracts were heated to 95 °C, cooled to room temperature, and treated with benzonase (Millipore, 70664) for 30 min at 37 °C. The benzonase treatment was repeated until the extract was free flowing. The protein concentration was determined and 50 μg protein aliquots were precipitated using acetone-ethanol. Precipitated protein was then digested with trypsin (1:100), once for 16 hours before another aliquot of trypsin is added (1:100) and incubated for an additional 4 hours. Peptides were then desalted using SepPak cartridges (Waters).

Peptides were analyzed by LC-MS/MS using a data-independent acquisition (DIA) approach implemented on a RSLCnano HPLC (Dionex) coupled to an Orbitrap Exploris 480 mass spectrometer (Thermo) using DIA windows reported previously. Peptides were separated on a 50-cm (2-μm particle size) EASY-Spray column (Thermo Fisher Scientific), which was assembled on an EASY-Spray source (Thermo Fisher Scientific) and operated constantly at 50□°C. Mobile phase A consisted of 0.1% formic acid in LC–MS-grade water and mobile phase B consisted of 80% acetonitrile and 0.1% formic acid. Peptides were loaded on to the column at a flow rate of 0.3□μl□min^− 1^ and eluted at a flow rate of 0.25□μl□min^− 1^ according to the following gradient: 2–40% mobile phase B in 120□min and then to 95% in 11□min. Mobile phase B was retained at 95% for 5□min and returned back to 2% a minute after until the end of the run (160□min in total).

The spray voltage was set at 2.2□kV and the ion capillary temperature at 280□°C. Survey scans were performed at 15,000 resolution, with a scan range of 350–1,500□*m*/*z*, maximum injection time 50□ms and AGC target 4.5□×□10^5^. MS/MS DIA was performed in the orbitrap at 30,000 resolution with a scan range of 200–2,000□*m*/*z*. The mass range was set to ‘normal’, the maximum injection time to 54□ms and the AGC target to 2.0□×□10^5^. An inclusion mass list with the corresponding isolation windows was used as previously reported. Data for both survey and MS/MS scans were acquired in profile mode. A blank sample (0.1% TFA, 80% MeCN, 1:1□v:v) was run between each sample to avoid carryover.

#### Proteomics Data Analysis

MS raw data files were processed using Spectronaut v.14.7.201007.47784 with a human reference FASTA sequence from UniProt, using default search parameters. The resulting protein-level data were analyzed using R v.3.5.0. Protein abundances are expressed as parts-per-million (ppm) and protein copies. To calculate ppm, protein intensities were divided by the total protein intensity and therefore reflect a concentration. To calculate protein copies, protein intensities were divided by the sum of histone intensities.

#### Metabolomics

Untargeted metabolomics analysis was performed by liquid chromatography-mass spectrometry (LC-Ms) on a Thermo Fusion interfaced with a Thermo UltiMate 3000 RSLC system. Samples were injected on to ZIC-pHILIC column (150 × 4.6 mm, 5μm; Merck SeQuant, Umea, Sweden) maintained at 40°C. Mobile phase A consisted of 20mM ammonium carbonate in water and mobile phase B consisted of acetonitrile. The percentage of mobile phase A was increased from 20% to 80% over 15 min and then to 95% held for 2 min before re-equilibration to the starting conditions over 9 min. The flow rate was 300 μL/min. The mass spectrometer was operated in polarity-switching mode over the mass to charge ratio (m/z) range 70-1000 at a resolution of 120,000. The raw LC-MS files were processed with IDEOM (Creek et al., 2012), which uses the XCMS (Smith et al., 2006) and mzMatch (Scheltema et al., 2011) software in the R environment. The annotation of metabolites was based on matching to the accurate mass of compounds in the HMDB, LIPID MAPS, MetaCyc and KEGG databases integrated in IDEOM together with comparison to an in-house authentic standard mix that was run alongside the samples. Lists of annotated metabolites and their relative abundances in the biological groups were retrieved and subjected to statistical analysis using MetaboAnalyst 5.0 (Pang et al., 2021).

## DATA AVAILABILITY

The mass spectrometry proteomics data have been deposited to the ProteomeXchange Consortium via the PRIDE partner repository with the dataset identifier PXD036519. Metabolomics data have been deposited to the EMBL-EBI MetaboLights database with the identifier MTBLS5868.

## ACKNOWLEDGEMENTS

We thank colleagues in the Ly, Saurin, and Barr groups for helpful discussions, Christos Spanos and Van Kelly (Wellcome Centre for Cell Biology) and Alex Montoya and Holger Kramer (MRC-LMS Proteomics core) for help with mass spectrometry analysis. Funding to TL was provided by the Wellcome Trust (206211/A/17/Z, 218305/Z/19/Z). Funding to ARB was provided by a CRUK Career Development Fellowship (C63833/A25729) and to RA and JAH through MRC-LMS core funding (MC-A658-5TY60). Funding to ATS was provided by a CRUK Programme Foundation Award (C47320/A21229), which also funded LC, and a Tenovus Scotland studentship, which funded RF. MB is supported by the Biotechnology and Biological Sciences Research Council EASTBIO DTP (BB/M010996/1). JAM is supported by a Medical Research Council Career Development Award (MR/M02122X/1) and is a Lister Institute Research Prize Fellow.

## FIGURE LEGENDS

**Supplementary Figure 1.**
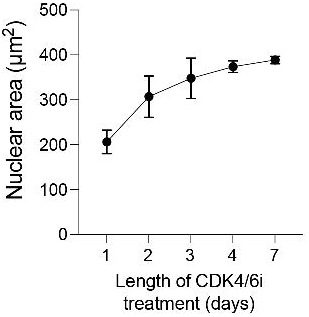
Nuclear area continues to increase during a prolonged CDK4/6i arrest. Mean +/- s.d. are shown.

**Supplementary Figure 2.**
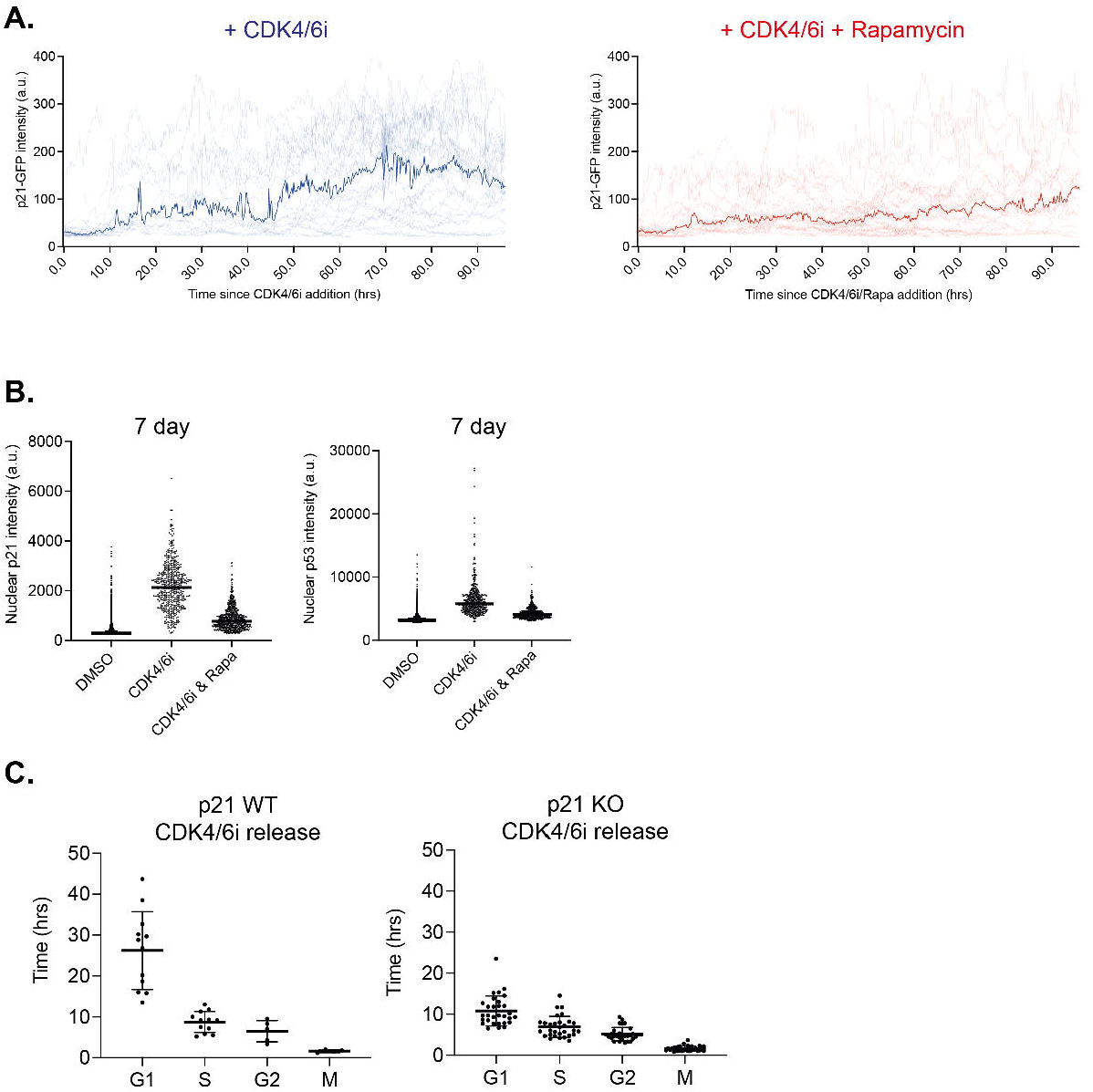
**A**. Graph showing single-cell data (faint traces) and median nuclear p21-GFP intensity (also plotted in Figure 2C) measured in RPE1 mRuby-PCNA p21-GFP H3.1-iRFP cells after addition of CDK4/6i (red curve) or CDK4/6i and rapamycin (blue curve). n=78 cells for CDK4/6i and n=66 cells for CDK4/6i and rapamycin. **B**. Graph shows nuclear levels of p21 and p53 protein, measured by immunostaining, in RPE1 cells treated for 7d with CDK4/6i or CDK4/6i and rapamycin. Each dot represents a single cell. **C**. Graphs show timings of cell cycle phases taken from Figure 2A. Only cell cycle phases up to the end of the first mitosis after inhibitor release and where the full phase was captured are shown.

**Supplementary Figure 3.**
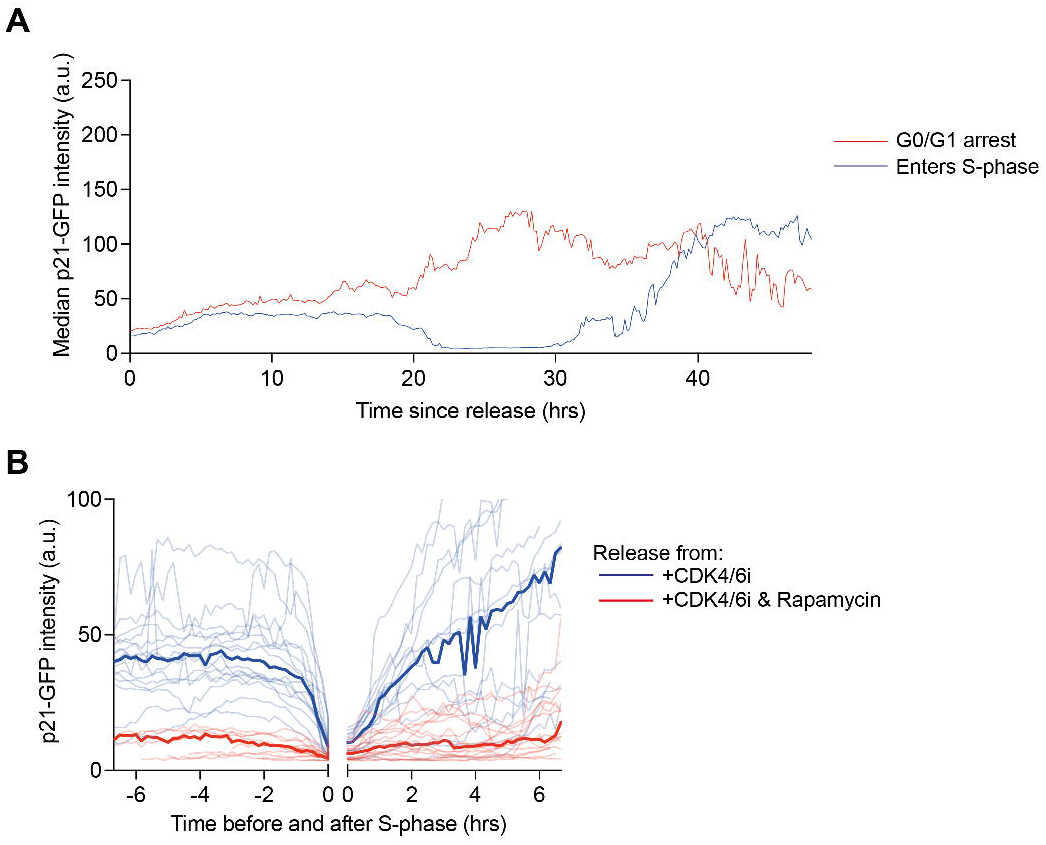
**A**. Median p21-GFP values from data shown in Figure 3A in cells released from CDK4/6i. Cells that remain arrested in G0/G1 are shown in red, cells that re-enter S-phase are shown in blue. **B**. Same data as shown in Figure 3A showing nuclear p21-GFP intensity in cells released from 7d CDK4/6i (blue curve) or CDK4/6i and rapamycin treatment (red curve). Here, only cells that enter and exit S-phase during the imaging period are shown and data are *in silico* aligned around S-phase entry and exit (time 0h). The line in bold is the median p21-GFP intensity and the faint lines are single-cell p21-GFP levels.

**Supplementary Figure 4.**
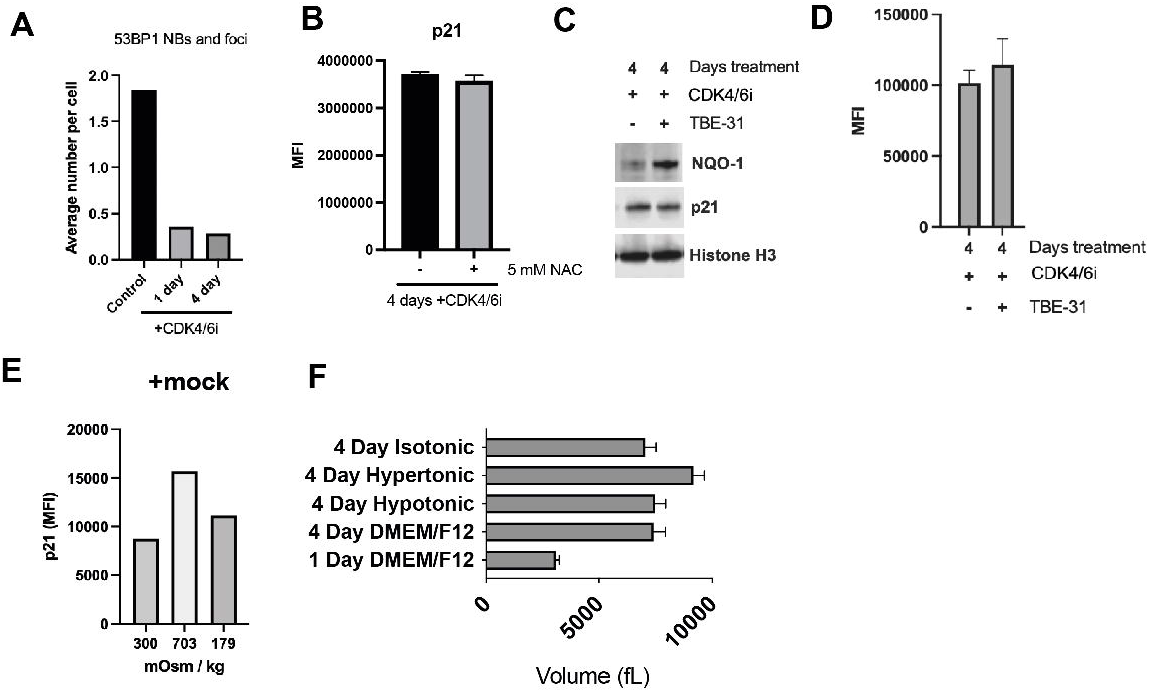
**A**. The frequency of 53BP1 nuclear bodies (NBs) and foci in control (DMSO-treated) versus CDK4/6i-treated cells. **B**. Flow cytometry analysis of cells co-treated with CDK4/6i for 4 days and -/+ 5 mM N-acetylcysteine (NAC) and immunostained for p21. MFI = geometric median fluorescence intensity. **C**. Immunoblot analysis of extracts from cells co-treated with CDK4/6i for 4 days, and either 100 nM NRF2 activator TBE-31 or vehicle control (acetonitrile). **D**. Flow cytometry analysis of cells treated identically to (C) and immunostained for p21. **E**. Flow cytometry analysis of asynchronous RPE1 cells treated with isotonic, hypertonic, or hypotonic media, and immunostained for p21. **F**. Measured volumes of cells treated with CDK4/6i and indicated media. Error bars indicate s.d. (n ≥ 90 cells).

**Supplementary Figure 5.**
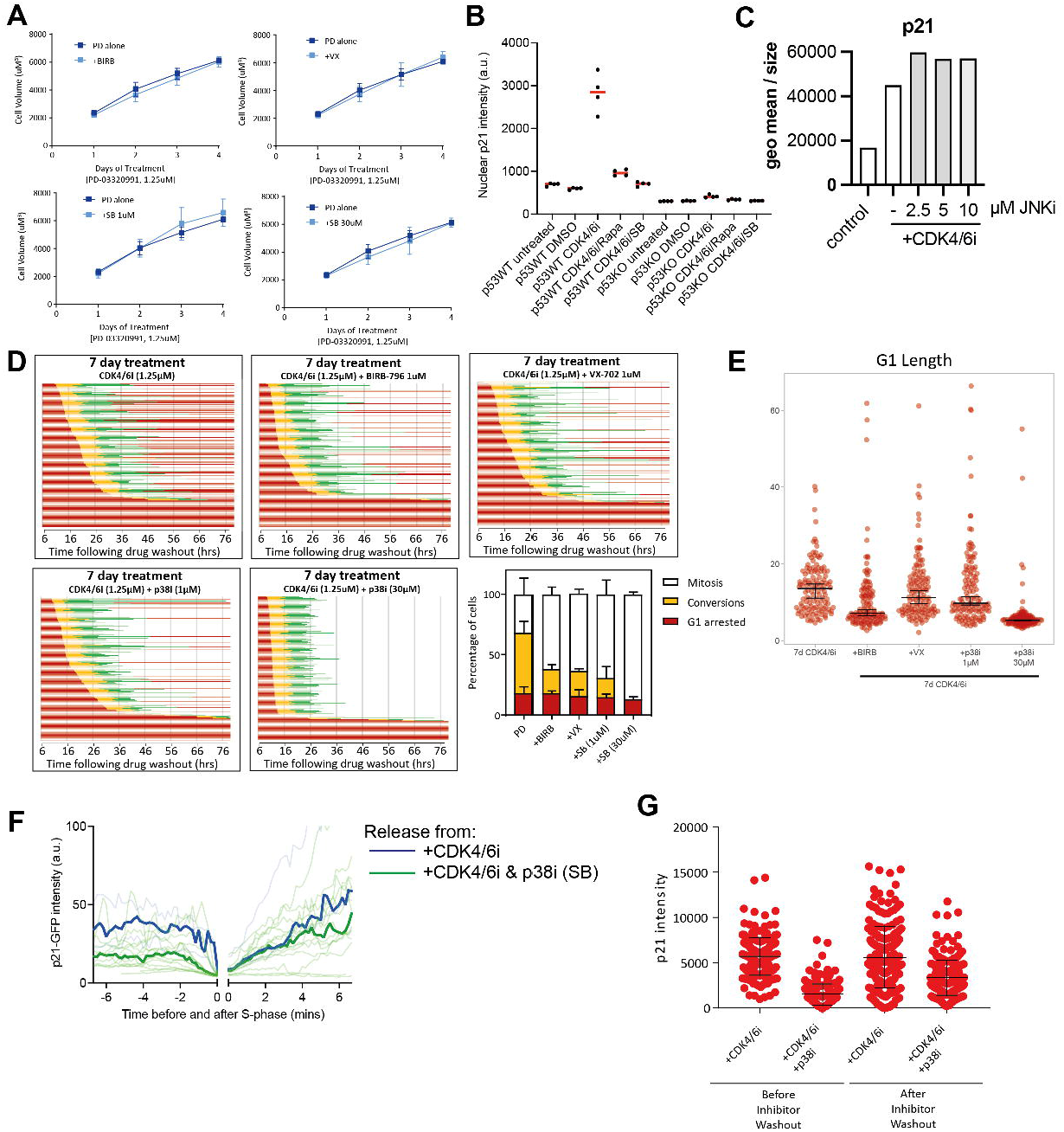
**A**. Graphs showing cell volume changes during incubation with CDK4/6i alone or combined with different p38i. n=3. Mean +/- s.d. are shown. **B**. Graph shows nuclear levels of p21 protein, measured by immunostaining, in RPE1 cells treated for 7d with CDK4/6i, CDK4/6i and rapamycin or CDK4/6i and p38i (30 μM SB-203580). Each dot represents a technical repeat. Red line is the mean. **C**. Flow cytometry of RPE1 cells immunostained for p21. Geometric mean of fluorescence intensity was normalized to size (forward scatter). Cells were co-treated with CDK4/6i and the indicated concentrations of JNK-IN-8 (or vehicle control) for 4 days. Control cells were untreated asynchronous cells. **D**. Graphs show FUCCI RPE1 cells released from 7d arrest with indicated inhibitors into the Eg5 inhibitor STLC to arrest cells in mitosis. Live cell imaging was started 6h after drug washout. Red is G1, yellow is S-phase, green is G2. Cells that withdraw from the cell cycle in G2 return to red colour as APC/C^Cdh1^ becomes active and the Geminin reporter is degraded. Insert graph shows mean of n=3 +/- s.d. **E**. Graph shows length of the first G1 after release from 7d treatment with inhibitors. Each dot represents a single cell. The horizontal lines display the median, and error bars show 95% confidence intervals. **F**. Same data as shown in Figure 5E but here only cells that entered S-phase are shown and tracks are aligned to S-phase exit. **G**. Quantification of p21 intensities in RPE1 cells treated with CDK4/6i + p38i (30 μM SB-203580). Cells were treated for 7d and p21 intensities were analysed by immunostaining before or after drug washout (48 h washout period). 100 cells were analysed. Mean of n=1 +/- s.d. between individual points.

**Supplementary Figure 6.**
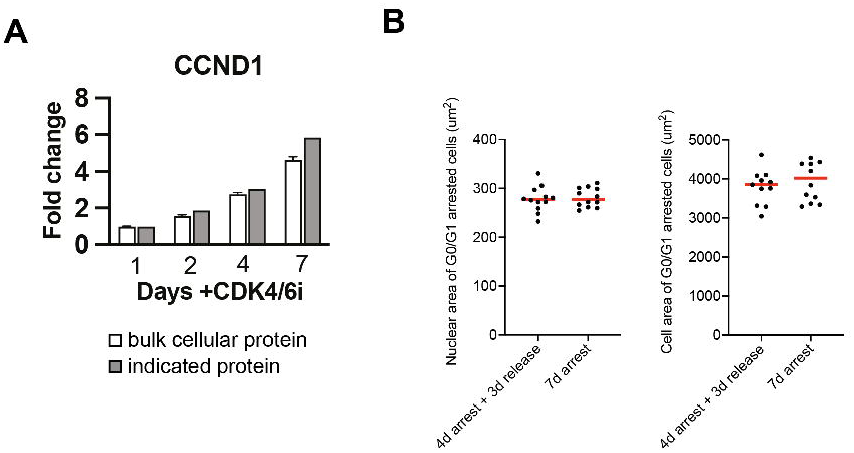
**A**. Graph shows data from quantitative proteomics that Cyclin D1 (CCND1) superscales during cellular overgrowth in CDK4/6i. **B**. Cells were arrested for 4d in CDK4/6i. CDK4/6i was then washed out into media containing EdU +/- CDK4/6i and cells fixed 3d later. Only EdU negative cells (G0/G1 arrested cells) were quantified here. Each dot represents a technical repeat. Red line is the mean.

## SUPPLEMENTARY TABLES

**Supplementary Table 1. Proteomics dataset**.

